# Force-insensitive myosin-I enhances endocytosis robustness through actin network-scale collective ratcheting

**DOI:** 10.1101/2025.04.04.647278

**Authors:** Michael A. Ferrin, Ross T.A. Pedersen, David G. Drubin, Matthew Akamatsu

**Affiliations:** Department of Molecular and Cell Biology, University of California, Berkeley, Berkeley, CA, USA; Department of Biology, University of Washington, Seattle, WA, USA

**Keywords:** Actin, Arp2/3 complex, Myosin-I, clathrin-mediated endocytosis, mathematical modeling

## Abstract

Force production by Type-I myosins influences endocytic progression in many cell types. Since different myosin-I isoforms exhibit distinct force-dependent kinetic properties, it is important to investigate how these properties affect endocytic outcomes, and the mechanisms through which myosin-I contributes to endocytosis. To this end, we adapted our agent-based simulations of endocytic actin networks and incorporated nonprocessive, single-headed myosin motors at the base of the endocytic pit. We varied the unbinding rate and the force dependence of myosin unbinding. Our results revealed that the inclusion of myosin motors facilitated endocytic internalization, but only under kinetic regimes with rapid and less force-sensitive unbinding. Conversely, slow or strongly force-dependent unbinding impeded endocytic progression. As membrane tension increased, the boundary between assistive and inhibitory phases shifted, allowing the myosins to assist over larger regions of the kinetic landscape. Myosin-I’s contribution to internalization could not be explained by direct force transduction or increased actin assembly. Instead, the myosins collectively bolstered the robustness of internalization by limiting pit retraction.

**Significance Statement:**

- Type-I myosins with varying force sensitivity levels participate in different membrane deformation pathways, but the mechanistic link between molecular biophysical properties and cellular function remains poorly understood.
- The authors analyze a computational model of endocytosis with type-I myosins and find that myosins with lower force sensitivity assist endocytosis by reducing backsliding along the internalization trajectory, while myosins with higher force sensitivity stall endocytosis by sequestering actin in non-productive orientations.
- These results introduce a new perspective on the function of type-I myosins in membrane reshaping: as a collective emergent property rather than the sum of individual force-generating motors.

## Introduction

Type I myosins are non-processive, membrane-associated molecular motors that generate a sheer force on actin filaments via a mechanochemical cycle that couples ATP hydrolysis to a “power stroke” conformational change [1]. Different myosin-I isoforms function in many distinct membrane trafficking pathways: vertebrate myosin-1b in the generation of post-Golgi carrier vesicles [2], myosin-1c in some examples of compensitory endocytosis [3], and vertebrate myosin-1e and its orthologs (including Myo5 in S. cerevisiae) in clathrin-mediated endocytosis (CME) [4–6]. Detailed single-molecule studies of several of these isoforms have revealed a rich diversity of mechanochemical properties [7–11]. One notably variable property among type I myosins is the force dependence of their detachment rate from actin filaments (Figure 1 A-B). Vertebrate myosin-1b is so acutely force-sensitive that it forms a “catch bond” that is more suited to function as a tension-sensitive anchor [9], while S. cerevisiae Myo5 has minimal force sensitivity and is better suited to function as a power-generating transporter [7].

**Figure 1.**
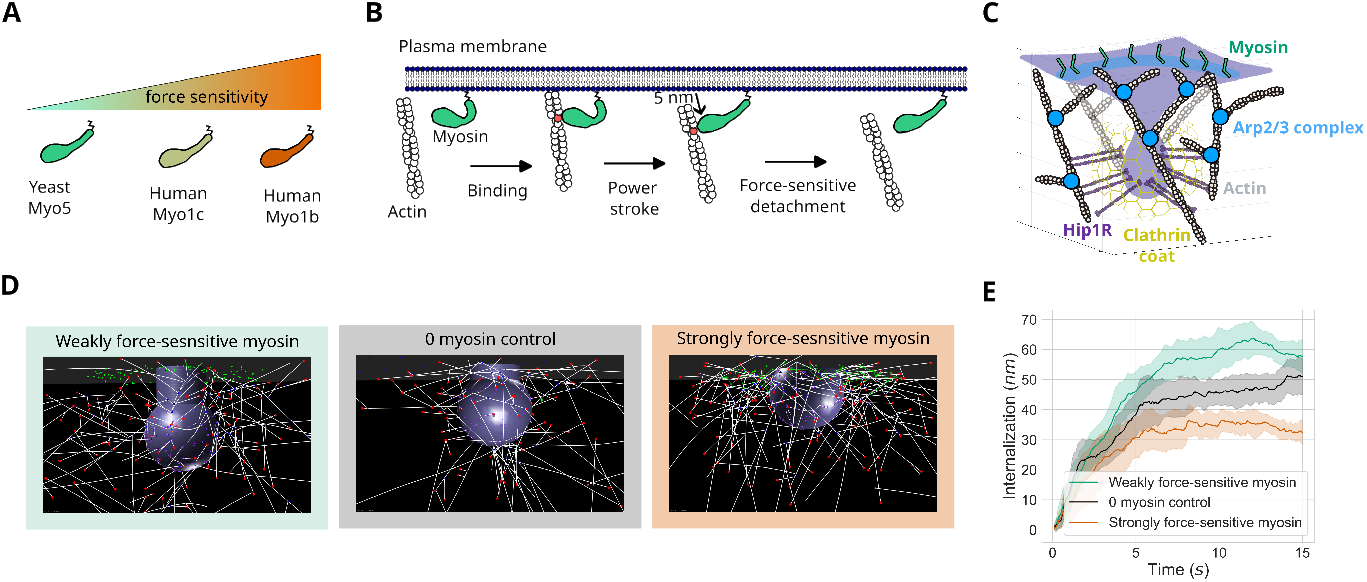
Weakly force-sensitive myosin-1 can assist endocytosis in computational simulations. (A) A diversity of actin unbinding kinetics exists among type-I myosins involved in different cellular processes, from the force-insensitive motor Myo5 to the force-sensitive anchor Myo1b. (B) Schematic of type-I myosin activity. Membrane-bound type I myosins can bind to actin filaments, take a single step, and then release the actin at a force-dependent rate. (C) Schematic of components included in agent-based model of clathrin-mediated endocytosis. (D) Snapshots of simulation endpoints in 3 different conditions: with weakly force-sensitive myosin (left), without myosin (center), with strongly force-sensitive myosin (right). (E) Pit internalization over simulated time for the three conditions in (C), averaged over 12 simulations each. Error bars are 95% confidence intervals using the t-statistic. All simulations were run with a resistance to pit internalization of 0.2 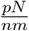.

Among all the intracellular behaviors associated with type I myosins, their involvement in CME has been examined in the most molecular detail. CME serves as the primary pathway for internalizing extracellular and plasma membrane-associated cargo in eukaryotic cells. As cargo adapters and curvature-generating proteins gather on the intracellular surface of a patch of the plasma membrane, the membrane invaginates into the cytoplasm and eventually pinches off to form a vesicle. While passive components that induce curvature upon assembly are adequate to transform a relaxed membrane into a vesicle [12, 13], additional active force from actin polymerization is essential to overcome resistance from membrane tension or turgor pressure [14–19]. Type I myosin additionally aids in actin-based force generation during CME, as its depletion completely halts endocytic internalization in budding yeast [14, 20–22] and partially disrupts internalization in mammalian cell culture [23]. Notably, the force-sensitive actin binding kinetics have been measured for the budding yeast endocytic myosin Myo5 [7], and homologous myosins in other simple eukaryotes [24–27] and vertebrates [28] have been characterized in detail the absence of resistive force. Given that clathrin-mediated endocytosis has been studied in rich quantitative detail, we utilized it as a model membrane-budding process to investigate how the various biochemical properties of type I myosins may influence their mechanistic role in membrane morphogenesis.

A minimal model of actin cytoskeletal assembly in CME has previously demonstrated that the spatial segregation of actin filament linker proteins on the invaginating membrane and branched actin nucleators around the pit’s base are sufficient to create a self-organizing actin network that generates an internalizing force [29]. However, this minimal network was incapable of internalizing the membrane to the degree predicted to be necessary for reaching the scission stage of CME. Given the assumed additional force-generating capacity of type I myosins, we inquired whether incorporating them could enhance internalization in this model. Additionally, we utilized the power of biophysical modeling to conduct *in silico* perturbations and take measurements that are not experimentally accessible to predict the design principles and mechanochemical parameters of a type I myosin optimally tuned for CME.

Simulations of our model across a variety of myosin parameters and mechanical environments revealed that the contribution of type I myosin to endocytic membrane internalization strongly relies on the force sensitivity of its actin-binding kinetics. While highly tension-sensitive tether myosins stall endocytosis, myosins that are relatively force-insensitive motors enhance average internalization. We find that tether myosins inhibit internalization by sequestering actin filaments in orientations that are not conducive to transmitting force to the membrane invagination. In contrast, motor myosins subtly influence actin network reorganization and limit pit retraction for more effective internalization.

## Results and Discussion

### Type-I myosin can assist internalization in CME simulations

To model the contribution of type I myosin to CME, we built on our model of branched actin assembly that generates force on an endocytic pit against membrane tension [29], utilizing the agent-based cytoskeletal modeling platform Cytosim [30]. Briefly, our model is initialized with the following components and rules. We represent the clathrin-coated pit and membrane neck as a solid sphere and a cylinder with a smaller radius than the sphere, respectively. Since the CME actin module functions following the initiation of membrane curvature development [31], we initialize our simulations with the pit partially internalized. The actin-binding endocytic coat linker protein (Hip1R) is randomly distributed over the surface of the pit. The plasma membrane base from which the pit internalizes is a flat surface, where pre-activated branched actin nucleators (Arp2/3 complexes) are arranged in a ring around the pit [32, 33]. A small number of actin mother filaments start with random positions and orientations near the pit, allowing them to diffuse according to 3D Brownian dynamics. When they come close to an active Arp2/3 complex at the membrane, they can spontaneously bind and nucleate a new daughter filament at a 70° angle [34, 35]. The filaments polymerize and stochastically undergo capping at experimentally estimated rates. Filaments can also stochastically bind to Hip1R when in close proximity. Mechanical force from the spring-like behavior of these transient bonds and collisions with the membrane, along with Brownian fluctuations, can bend an actin filament according to its persistence length [36]. Informed by membrane mechanics simulations, we model membrane tension as spring-like resistance of the endocytic pit, with a linear relationship between internalization distance and the force required to maintain it there. As previously shown [29], these components alone are sufficient to reproducibly drive internalization up to a stall distance of ~50 nm (Figure 1 D-E).

In the present work, we added type I myosins to the base membrane in a ring around the pit (Figure 1 C) [32, 37]. These myosins can stochastically bind to actin filaments, upon which they exert a power stroke to step up to 5 nm toward the barbed end of the filament [7] and then unbind (Figure 1 B). To model a catch bond, the myosin’s unbinding rate decreases with increasing applied force of the myosin-actin bond [11] (Figure 1 B). We were able to constrain the majority of the model parameters to available experimental measurements and empirically fit the remainder to recapitulate the behavior of endocytic type I myosins in reconstitution experiments (Table S1, Figure S2).

Simulating this minimal CME model with type-I myosins showed that the modulation of pit internalization depended on myosin parameters (Figure 1 D-E). With highly force-sensitive myosins present, internalization was significantly hindered compared to simulations that omitted myosins under the same conditions (Figure 1 E). Conversely, including mildly force-sensitive myosins led to greater internalization than in simulations without myosin (Figure 1 E). After observing this relationship between myosin’s kinetic parameters and endocytic efficiency, we aimed to systematically investigate how the myosin’s force sensitivity parameter impacts endocytic outcomes.

### The kinetics of myosin’s force-sensitive unbinding determine its assistive capacity during endocytic pit internalization

Force-sensitive detachment kinetics are commonly modeled with the Bell Equation [38] (schematized in Figure 2 A and plotted in Figure 2 B):

**Figure 2.**
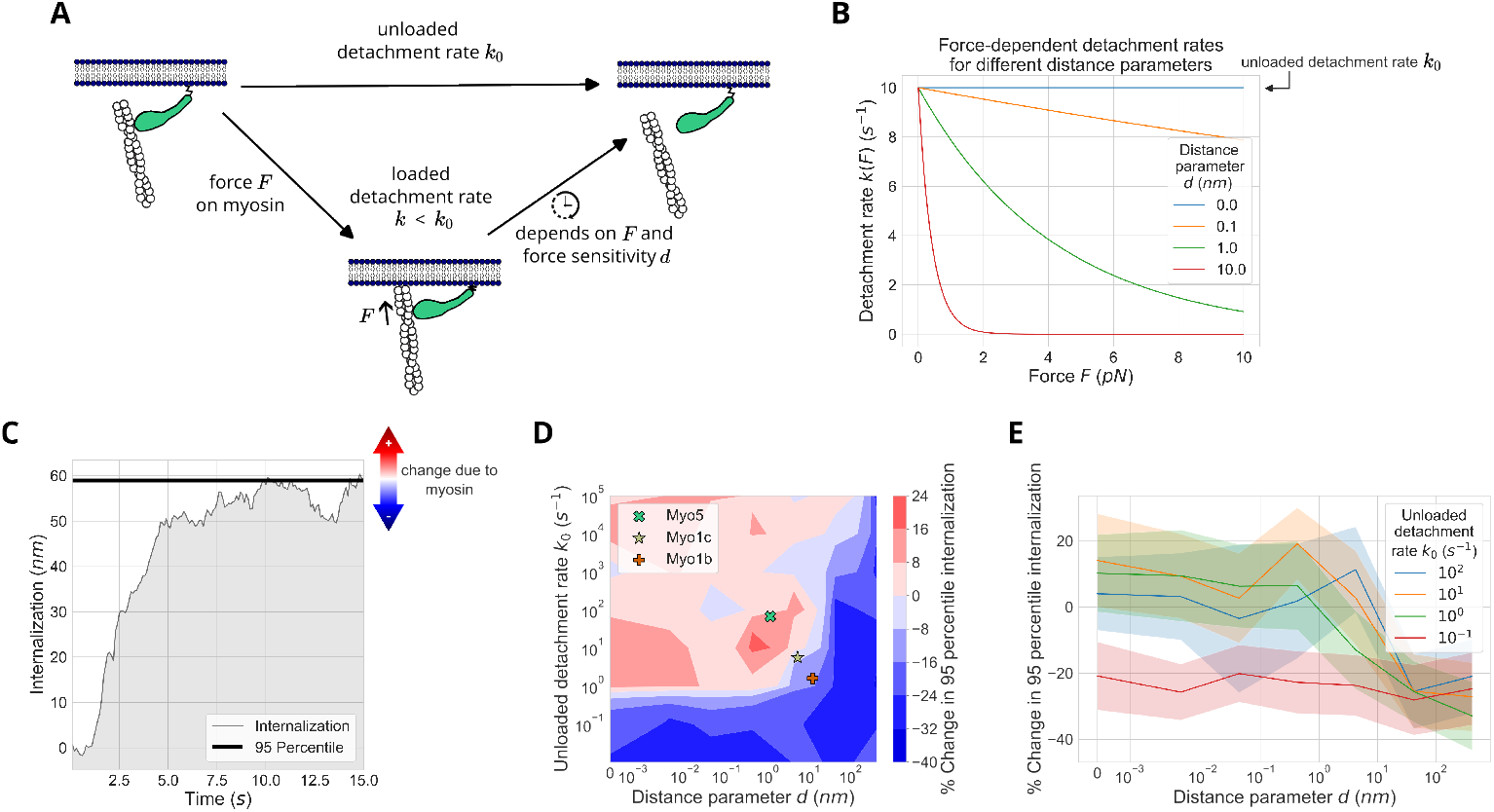
Kinetics of myosin’s force-sensitive unbinding determine its assistive vs. resistive capacity in endocytosis. (A) Cartoon schematic highlighting the relevant parameters for force-sensitive unbinding. The unloaded detachment rate *k*_0_ is defined as myosin’s detachment rate from actin filaments in the absence of applied force *F*. With applied force, the predicted detachment rate becomes a function of *k*_0_, *F*, and the distance parameter *d*. (B) Plots of the Bell equation provide a theoretical prediction of how the actin-myosin detachment rate *k*(*F*) responds to applied force *F* at different values of the distance parameter *d*. (C) Demonstration of the 95th percentile of internalization relative to the plot of internalization over time for a single simulation. This serves as a summary statistic for comparing many simulations. (D) Contour plot of the % change in mean 95th percentile of internalization relative to simulations with 0 myosins, over a range of parameter values for *d* and *k*_0_. At low *d* and high *k*_0_, myosins cause an increase in internalization, while they cause a decrease in internalization at high *d* and low *k*_0_. Type I Myosins with experimentally measured values [7, 9, 10] are mapped in the plot. (E) Plots with 95% confidence interval of the % change in mean 95th percentile of internalization over *d* for selected values of *k*_0_ (different lines). All simulations were run with a resistance to pit internalization of 0.2 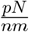.

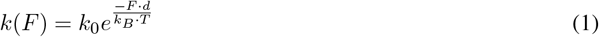

where *k*(*F*) is the effective detachment rate at applied force *F*, *k*_0_ is the unloaded detachment rate, *d* is the distance parameter characterizing force sensitivity, *k*_*B*_ is Boltzmann’s constant, and *T* is temperature. The outcome of varying distance parameter *d* can be visualized in Figure 2 B: the detachment rate is completely force-insensitive at *d* = 0, and the detachment rate becomes more dependent on applied force *F* as *d* increases.

We use the 95th percentile statistic of internalization as a single value summarizing endocytic success for comparison across various conditions (Figure 2 C; see methods). Visualizing the percentage change in the 95th percentile of internalization relative to simulations with no myosins revealed clear boundaries between myosin kinetic parameters that facilitate endocytosis and those that inhibit it (Figure 2 D). We observed that internalization, on average, increased with myosins that have fast unloaded detachment (high *k*_0_) and weak force sensitivity (low *d*) (Figure 2 D). Notably, overlaying the single-molecule measurements of these parameters for known type I myosins [7, 9, 10] demonstrated that the kinetic parameters for Myo5, a myosin involved in CME, lie in the facilitative regime, while the two type I myosins unrelated to CME, Myo1c and Myo1b, fall within the non-facilitative regime (Figure 2 D). Accounting for variability between simulations, the distinction between assistive and stall-inducing regimes was sharpest between *d* of 5 and 50 nm, for most values of *k*_0_ (Figure 2 E).

### The mechanical environment influences the optimal myosin parameters for endocytosis

The necessity for actin-based force generation in CME strongly depends on the magnitude of forces opposing internalization: while impairing actin assembly has a minimal effect on endocytic success in mammalian cells with low membrane tension, the same perturbation almost completely stalls internalization when membrane tension is high [18]. Similarly, reducing osmotic pressure can even partially relieve the requirement for actin assembly in budding yeast CME [16]. Given that the depletion of Myo1e from mammalian cells results in similar endocytic defects as actin drug treatment [23], we hypothesized that the assistive role of of myosin-I in CME should depend on the magnitude of mechanical resistance from membrane tension or pressure in a manner qualitatively mirroring that of actin (Figure 3 A).

**Figure 3.**
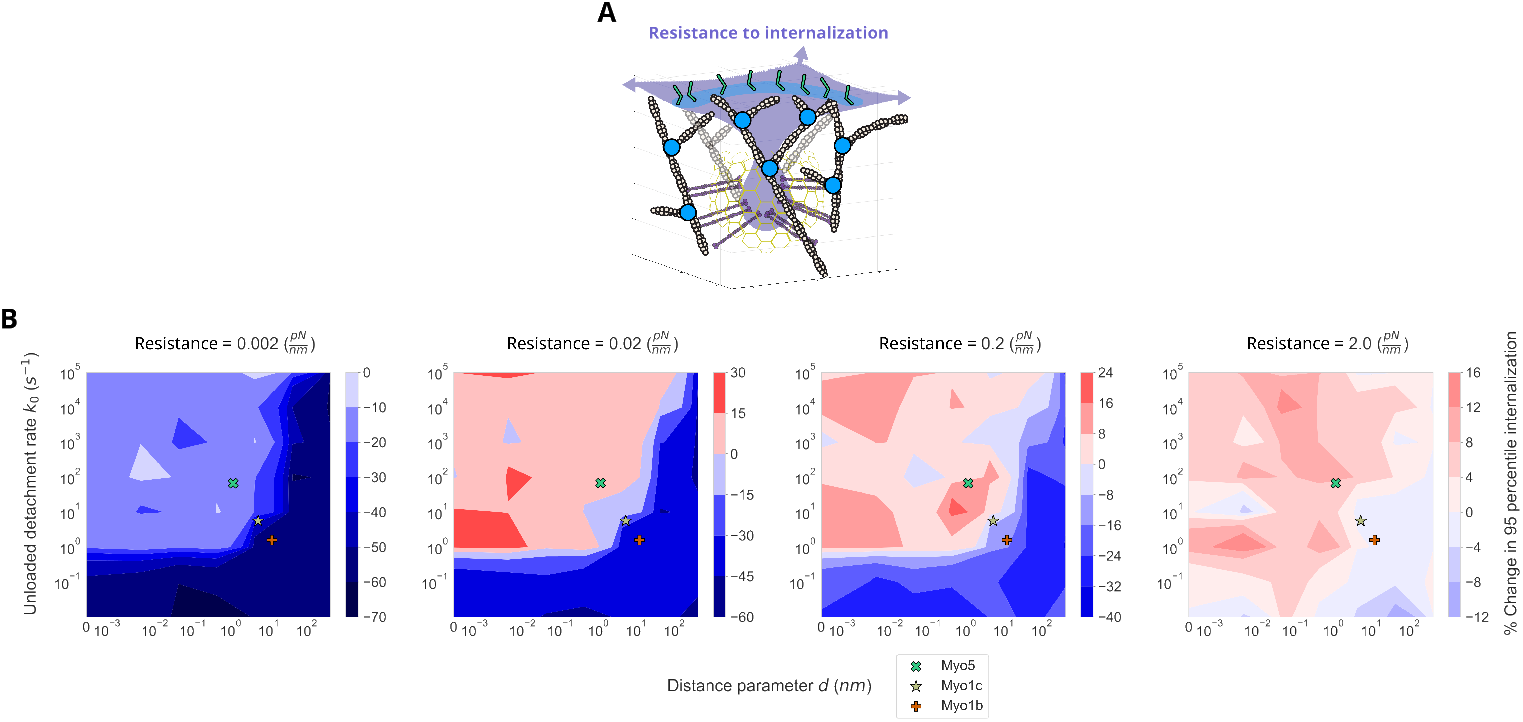
The mechanical environment influences the optimal myosin parameters for endocytosis. (A) These simulations explore how myosin’s influence on CME depends on the magnitude of force resisting internalization, which could arise from membrane tension or pressure. (B) We varied myosin’s distance parameter *d* and unloaded detachment rate *k*_0_ for simulations with varying resistance to internalization. The myosins were universally non-assistive to endocytosis at very low resistance, while they exhibited a boundary at intermediate resistance that levels out at high resistance.

To investigate for this yet experimentally unexplored impact of myosin under varying opposing force, we ran simulations with the same combinations of force-dependent kinetic parameters as described above, across a wide range of internalization resistances (Figure 3 B). We found that, under increasing values of membrane tension, the boundary between assistive and inhibitory contributions from myosin lessened and shifted, such that myosin became assistive over larger ranges of kinetic parameters. Although the size of the myosin-assistive regime increased with increasing membrane tension, the magnitude of assistance generally decreased.

This increase in the myosin-assistance regime with internalization resistance can be mapped to experimental measurements of membrane tension and myosin-I endocytic phenotypes. Previous theoretical analysis demonstrated linear scaling between internalization resistance of 0.6 to 1.6 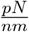 to plasma membrane tension of 0.15 to 0.45 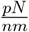 [29]. AFM measurements of SK-MEL-2 cells in isotonic media estimated a plasma membrane tension of 0.14 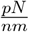 (corresponding to a resistance of ~0.6 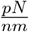 in our model), while hypotonic shock could elevate the tension to 0.29 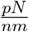 [17] (corresponding to a resistance of ~1.0 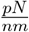). As our simulations span this range, these results are useful for predicting the yet unmeasured kinetic parameters of the mammalian endocytic Myo1e for maximizing robustness of CME. Interestingly, a myosin-I with force sensitivity and unloaded detachment rates corresponding to those of the budding yeast endocytic Myo5 is assistive to simulated internalization throughout 3 orders of magnitude around the resistance closest to a “relaxed” mammalian plasma membrane tension (Figure 3 B, 0.2 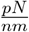). Yet other experimentally characterized non-CME type 1 myosins inhibit internalization at all resistance values, demonstrating that CME requires a fundamentally different myosin activity than the myosins that function in tension-sensitive anchoring can provide. Myosins with any parameters universally inhibit internalization at the lowest resistance, which is lower than the estimated tension for any cells measured to date [39].

### Myosins provide resilience against pit retraction

In seeking a mechanistic explanation for myosin’s binding kinetics-dependent function in endocytic outcomes, we observed that assistive myosins reduced pit retraction events at the nanometer scale. An inspection of individual internalization trajectories revealed that the pit did not steadily internalize throughout the simulation; rather, it transitioned between phases of internalization and retraction (Figure 4 A). Unlike the ¿100 nm-scale retraction events observed experimentally for endocytic sites lacking scission machinery [40], these sub-10 nm-scale retractions would fall below the resolution limit of time-lapse fluorescence microscopy. We investigated whether myosins modulate these retractions by quantitatively classifying retraction events and measuring their durations and distances. By comparing these retraction metrics across various myosin-I kinetic parameters, we identified a boundary of myosin parameters that closely mirrored our previous internalization measurements (Figure 4 B-C, compare with Figure 2 D). In the low *d* (weak force sensitivity) and high *k*_0_ (fast unloaded detachment) regime, where the 95th percentile of internalization was higher than in no-myosin controls (Figure 2 D), endocytic events spent less time in the retraction phase per simulation (Figure 4 B) and retracted shorter distances overall per simulation (Figure 4 C). These results suggested a mechanistic link between myosin’s control over pit retractions and endocytic outcomes.

**Figure 4.**
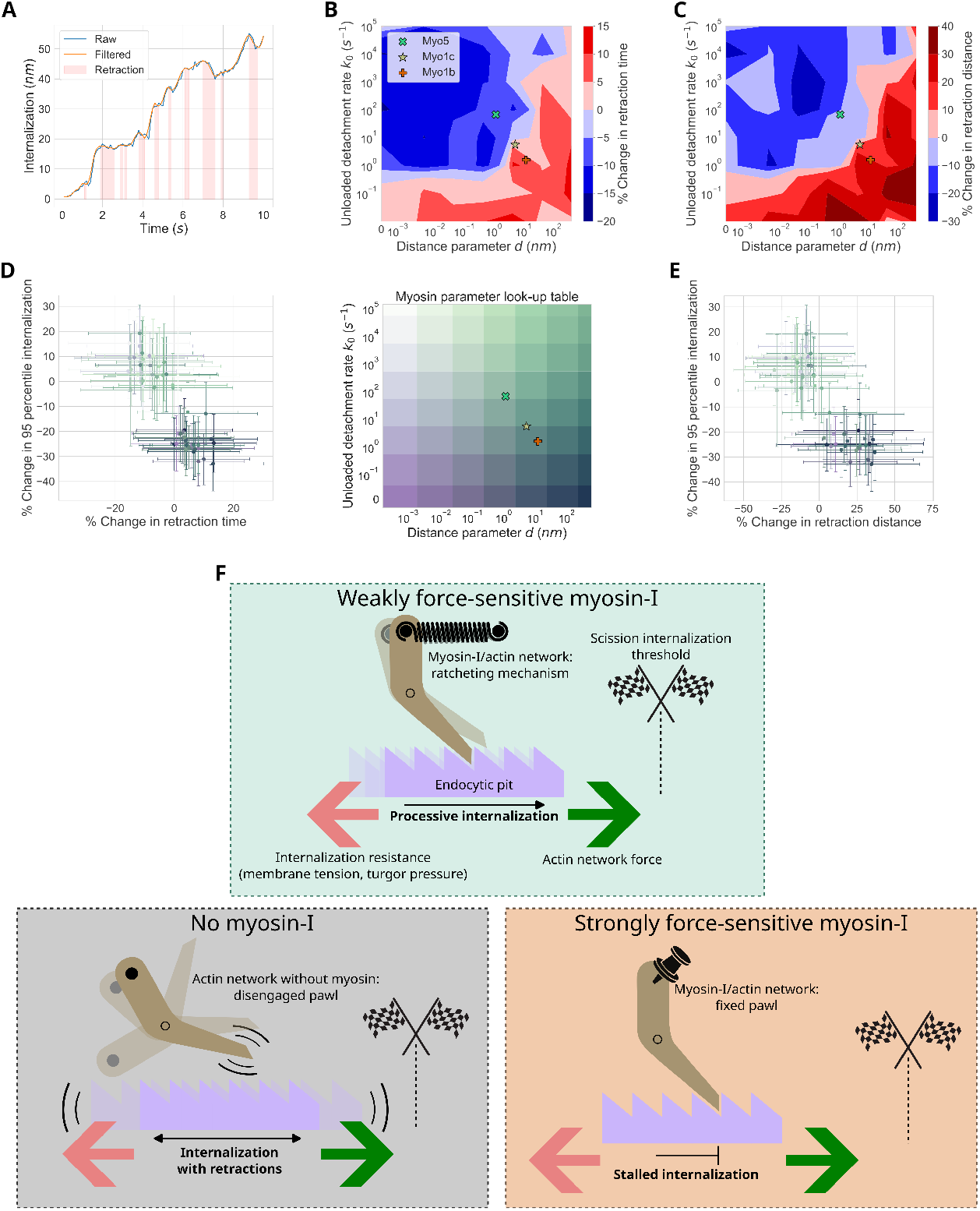
Assistive capacity of myosins for endocytic internalization can be explained by robustness to retraction. (A) Plot of raw and filtered internalization over time for a single simulation with retraction events highlighted in red (see methods for details). We analyzed retraction events in the first 10 simulated seconds of simulations. (B-C) Change in (B) mean total retraction time and (C) total retraction distance relative to no-myosin simulations. Both values were higher with high *d* (strongly force-sensitive) and low *k*_0_ (low unbinding rate) myosins, while the retraction measurements were lower with low *d* and high *k*_0_. (D-E) Change in mean 95th percentile of internalization relative to no myosins (y-axis) as a function of the change in mean total retraction time (D) and total retraction distance (E) (x-axes). Both metrics inversely correlated with change in internalization. We used a two-dimensional lookup table of myosin kinetic parameters to color code the data points. Each point with error bars represents the mean and 95% confidence interval of all replicate simulations of the same myosin parameter combinations indicated by color from the lookup table. (F) Cartoon conceptual model of type-I myosin function in CME. Endocytic internalization is like a ratchet with force from actin assembly pulling the ratchet toward the right (pit internalization). The kinetic parameters of myosin are analogous to the behavior of the pawl (brown). (Top) Myosins in the assistive parameter regime collectively behave as a balanced pawl spring, flexible enough to allow forward movement but strong enough to limit backsliding. (Bottom left) In the absence of myosin, the pawl spring is removed, allowing back-sliding of the ratchet. (Bottom right) If *d* is too high and *k*_0_ too low, the myosins behave like an immobilized pawl, stalling movement in any direction. All simulations were run with a resistance to pit internalization of 0.2 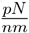.

To relate the effects of myosin-I on pit retractions to net internalization more quantitatively, we plotted the percentage change in the 95th percentile of internalization as a function of retraction time (Figure 4 D) and distance (Figure 4 E) for each myosin parameter combination. We identified a clear trend in which myosins that reduced retractions increased net internalization (Figure 4 D-E). In contrast, there was no apparent correlation between 95th percentile of internalization and retraction time (Figure S3 A) and distance (Figure S3 B) in simulations with no myosins included. Therefore, myosin’s influence on overall internalization may be explained by its ability to limit pit retraction.

These results lead us to a mesoscale conceptual model in which myosins modulate the ratchet-like behavior of the overall endocytic pit-actin system (Figure 4 F). CME can be envisioned as a ratchet driven by actin forces. Movement in the direction of the actin-based force is countered by membrane tension. In the presence of weakly force-sensitive myosins (Figure 4 F, top), the actomyosin network collectively acts like a spring-loaded pawl that enables the ratchet to be pulled by actin assembly in the membrane internalization direction while blocking retraction. In the absence of these myosins (Figure 4 F, bottom left), no equivalent spring exists to keep the pawl engaged with the ratchet, allowing the actin force to pull it toward internalization but making it susceptible to retractions due to fluctuations in the system. With strongly force-sensitive myosins (Figure 4 G, bottom right), the actomyosin network effectively pins the pawl, resisting any displacement of the ratchet. This “collective ratchet,” at the scale of the cytoskeletal network, differs from the Brownian ratchet mechanisms described for single molecules [41–43]. On a detailed molecular level, this ratcheting action may originate from myosin’s effect on actin rearrangement.

As the system’s geometry continually evolves over time with pit internalization, the actin network must continually reorganize to sustain a pulling force. Because of the stochastic nature of filament movements, this self-organizing network inevitably transitions through intermediate structures that do not maintain the same internalization force, leading to pit retractions. The weak force-sensitivity of myosin bonds might bias the actin network against rearranging in ways that hinder force production, thus reducing retractions. Indeed, we noticed subtle variations in individual actin filament organization due to the supportive myosins, such as an increase in radial distance from the pit and a flatter angle of actin barbed ends relative to the base membrane (Figure S4). This organization may indicate a mechanism that allows shallower filament angles to enhance forces from actin polymerization [44]. Future modeling and experimental efforts are needed to explore myosin’s impact on actin reorganization more fully.

Importantly, many outcomes of these simulations have also been observed experimentally. *In vitro* bead motility assays with branched actin assembly showed that Drosophila Myo1d inhibited bead motility in some conditions, including high filament capping, high concentrations of myosin, or “rigor” myosins that cannot unbind actin [45]. Our simulations recapitulated this inhibition in conditions of little or no myosin unbinding (Figure 2 D). In experiments, the presence of myosin led to a decrease in actin assembly by 20-30%, which was recapitulated quantitatively by *in silico* bead motility simulations as well as our endocytic actin simulations [45], (Figure S5 A-C). Each of these similarities with experimental observations help to validate the model.

Several other mechanisms for the function of myosin-Is in force-generating branched actin networks have been proposed, but these were not supported by our observations. Correlations between the presence of myosin motors, the size of endocytic actin networks, and pit internalization rates in live budding yeast suggest a model in which myosins assist CME by pulling actin filament barbed ends away from the plasma membrane, thereby reducing the “stall” force on barbed ends that would otherwise limit filament elongation [46]. Single-molecule characterization of Myo5’s binding kinetics indicates that myosin assists CME by directly applying force to actin filaments [7]. The aforementioned joint biochemical reconstitution and computational modeling study proposes a conceptually similar model in which myosins enhance branched actin force production by pushing actin filaments away from the membrane surface, strengthening the network by modulating the branching frequency along filament length [45]. However, we observed minimal changes to actin filament elongation rates due to the presence of myosin in the model (Figure S5 D). Furthermore, we found that myosin’s assistance to internalization persisted even without motor activity (Figure S6 A-B).

It is important to note that our model omits several microscopic properties of myosin that may account for some of the discrepancies between our simulation results and previously proposed mechanistic models. These omissions include endocytic myosins’ nucleation promoting factor (NPF) activity [14, 47], recruitment of downstream NPF proteins [23], the torque on actin resulting from myosin’s power stroke [48], a “failure force” that disrupts myosinactin bonds, and the capacity of myosins to reversibly attach to and diffuse along the membrane. We anticipate that each of these features will be systematically incorporated into future computational models and assessed for their emergent influence on the entire system, much like this work expands on an earlier CME model that lacked any myosins [29].

## Acknowledgments

We thank Sam Smith, Leanna Owen, Jennifer Hill, Tom Pollard, and members of the Drubin/Barnes laboratory for helpful discussions. We thank the Berkeley Reserach Computing team for maintaining the computing resources essential to carrying out this work and for technical support. This work was supported by the NSF Graduate Research Fellowship Program to M.A.F, as well as NIH grants NIGMS F32GM142145 to R.T.A.P., R35GM118149 to D.G.D., and NIGMS R00GM132551 to M.A. The content is solely the responsibility of the authors and does not necessarily represent the official views of the NIH.

## Materials and methods

All modeling and simulation code is available at: https://github.com/DrubinBarnes/ferrin_cme_myosin_cytosim

### Cytosim model

We adapted the agent-based model developed in [29] to run 3D stochastic simulations of endocytic pit internalization via a minimal branched actin network. In short, we model the endocytic pit as a solid sphere connected to a flat membrane base by a solid cylindrical neck. The pit experiences a spring-like resistance to internalization set by the parameter “bud confine.” Actin filament-binding linker proteins are randomly distributed over the surface of the pit. Pre-activated branched actin nucleators are randomly distributed in a ring around the pit, where they can freely diffuse along the flat membrane surface until they bind actin filaments. Myosins, when included, are also randomly distributed in a ring around the pit, though they cannot diffuse along the membrane. To initialize the event with a small number of actin filaments, linear actin filament nucleators are randomly distributed in a ring around the base of the pit, and subsequently are free to diffuse throughout the cytoplasm. Filaments can elongate, become stochastically capped, bind linker proteins emanating from the surface of the membrane at the nascent pit, bind branch nucleators to nucleate daughter filaments, and bind to myosins. Filaments also have a bending rigidity, so they store and release mechanical energy upon bending and straightening, respectively. The forces from actin polymerization against the membrane while bound to membrane-associated linker proteins is sufficient to internalize the pit 50-60 nm against the resisting force [29].

For main results, CME simulations were run for 15 simulated seconds, with simulation steps every 0.00005 s, and outputs reported every 0.1 s. Most model configurations analyzed in this work were run in replicates of 12, though simulations that terminated prematurely due to numerical instabilities were excluded from all analyses. See Figure S1 for a comprehensive replicate count for each configuration. Simulations for validating myosin non-processivity (Figure S2 B) and force-dependent attachment lifetime (Figure S2 C) were run for 1 simulated second with outputs reported at every 0.00005 s simulation step. Except where explicitly noted, we ran CME simulations with the pit resistance to internalization set at 0.2 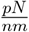.

The modifications made from [29] were changing the Arp2/3 branching angle from 77° to 70° to better fit recent experimental evidence [49] and adding myosin-I motors, as detailed below.

For a full list of parameters used in the model, see Table S1.

#### 0.0.1 Myosin-I model and parameter choices

Because the default model motor proteins defined in Cytosim are processive, i.e., they continuously displace along filaments while attached, we developed a new non-processive motor class for our model. To create the new motor, we modified the “mighty” class of Cytosim’s source code, which is a copy of the “motor” class allocated for such customized functionalities. This motor stops moving along the filament and remains attached after its initial displacement along the filament, thereby modeling a single power stroke for a single-headed myosin (Figure S2 A, B). We modeled myosins with abolished motor activity (Figure S6 A) by setting the displacement speed to 0.

The force-sensitive ADP release and ATP binding transitions of type I myosin’s ATPase cycle result in its force-sensitive detachment property [11] (Figure 1 B). We implemented this force-sensitive detachment by inter-converting between the Bell equation (Equation 1) and Kramers’ theory used in Cytosim (Equation 2):

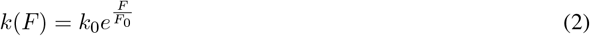

where *k*(*F*), *F*, and *k*_0_ are defined the same as in Equation 1, and *F*_0_ is the “unbinding force” parameter in Cytosim. By setting the remaining quantity from Equation 1 *T* = 303.15 *K*, we fully defined the mathematical relationship to convert between myosin distance parameter (*d*) and unbinding force (*F*_0_). The unloaded detachment rate (*k*_0_) and distance parameter (*d*) of the Bell equation (Equation 1) can be fit from single-molecule optical trapping experiments, thus providing a direct map between simulations and experiments. To validate this implementation, we simulated 2D filament gliding motility powered by myosin motors with parameters corresponding to measurements from the budding yeast endocytic Myo5. The resultant simulations qualitatively recapitulated its experimentally observed force-dependent attachment lifetime (Figure S2 C) and gliding actin motility rates (Figure S2 D) [50].

We set the parameters of myosin-I that were not the primary subject of investigation in this work based on experimental measurements of the budding yeast endocytic Myo5 where possible (Table S1). The actin-binding rate of 3 *s*^*−*1^ comes from the steady-state ATPase rate of phosphorylated Myo5 in saturating ATP, which measures the rate-limiting step of the Myo5 ATPase cycle for actin binding [50]. Myosin’s “max speed” sets the step size of a single power stroke, which we fit to 5 nm for Myo5’s total step size per ATPase cycle [50] (Figure S2 B). Except where explicitly noted, we set the baseline number of myosin-I molecules at 100, based on estimations of the total number of budding yeast type-I myosins recruited per CME site from live cell fluorescence microscopy [46, 51]. We set the radial distance of myosins around the pit based on super-resolution microscopy from budding yeast [32] and correlative super-resolution light and transmission electron microscopy in HeLa cells [37].

For other myosin-I simulation parameters that could not be directly constrained by experimental measurements, we empirically determined appropriate values by performing parameter sensitivity analyses. While the myosin-I motor domain is connected to the membrane through a lever arm and tail domain, the Cytosim model myosin is only comprised of the motor domain that cannot move in space. In order to model this spatial separation between the motor domain and the membrane, we placed the myosins at a fixed offset distance from the membrane (Figure S2 A). Inspection of myosin-I structural models informed our hypothesized 8 nm typical offset distance [48], and parameter scans revealed no significant impact on internalization around this range (Figure S2 E). The model myosin has a binding range, which is the maximum distance between the myosin and actin in which a bond can be established (Figure S2 A). Though this quantity may be related to myosin’s radius of gyration around its membrane-immobilized tail domain, there is no obvious direct physical complement to this parameter. We hypothesized a binding range of 4 nm, and found no significant impact on internalization around this range (Figure S2 F). Molecular bonds in Cytosim are modeled as springs with a stiffness parameter (Figure S2 A). As there is no direct physical correlate to this parameter of myosin, we scanned several orders of magnitude and chose a value of 100 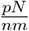 (Figure S2 G). Below this value, bonds stretched physically unrealistic distances *>*10 nm in CME simulations. Above this value, internalization was severely hindered for myosins with parameters tuned to Myo5 (Figure S2 G). Myosins also have a stall force, which is the force required to decrease their step size (Figure S2 A). This has not been measured for Myo5, so we picked an arbitrarily high value in order to guarantee stepping and subsequent unbinding in our simulations. The greatest pit internalization occurred with a high stall force for force-insensitive myosins, but we observed no significant trend for Myo5-tuned myosins (Figure S2 H).

### Simulation analysis

We wrote and modified custom Python scripts to parse simulation outputs, analyze results, and generate plots. Major concepts used in analysis are explained below.

#### 0.0.2 95th percentile internalization

We use the 95th percentile internalization as a metric summarizing the extent of pit internalization for a single simulation. This value, less sensitive to noise than the final time point or maximum internalization, is defined as the 95th percentile of internalization values at all time points throughout the 15-second simulation.

#### 0.0.3 Statistics summarizing differences relative to no myosin

In order to quantify the effect of a given myosin-I configuration on a simulation summary measurement (such as 95th percentile internalization), we calculate the % change in the mean measurement relative to no myosins with the function *f* (Equation 3):

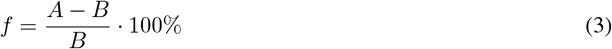

where *A* is the mean of the measurement in replicate simulations with myosin and *B* is the mean of the measurement in replicate simulations without myosin.

We propagate the uncertainty of the two means in the function by calculating the standard deviation *σ*_*f*_ Equation 4:

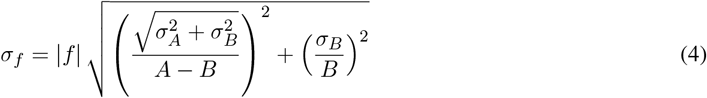

in which *σ*_*A*_ is the standard deviation of the measurement in replicate simulations with myosin and *σ*_*B*_ is the standard deviation of the measurement in replicate simulations without myosin.

We can then calculate the 95% confidence interval CI with Equation 5:

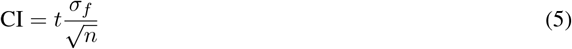

where *t* is the t-statistic for a 95% confidence interval (two-tailed) and *n* is the minimum number of replicates between the with- or without-myosin measurement.

#### 0.0.4 Internalization retraction analysis

We carried out the following steps to measure internalization retraction time and distance per simulation:

1. We applied a Savitzky–Golay convolution filter to smooth each internalization trajectory. We used a window size of 10 time points (1 second simulated time) and a 4th-order polynomial for fitting.
2. We used a peak-finding algorithm to identify all local minima and maxima in the smoothed trajectory. The duration between a local maximum and the next local minimum is defined as a retraction event. The length of this duration is the retraction time, and the difference in smoothed internalization between the start and end of the retraction event is the retraction distance.
3. We summed the retraction times and distances for all retraction events in the first 10 seconds of the simulation to arrive at the final measurements per simulation.

#### 0.0.5 2D Lookup table

To create a quantitative colormap that represents both myosin parameters *d* and *k*_0_, we created a 2D lookup table that varies two color axes corresponding to different combinations of myosin parameters.

#### 0.0.6 Myosin binding event analyses

The binding state of each individual myosin was tracked at each simulation time point to identify all binding events. The information associated with the actin filament participating in each binding event was used for subsequent analyses.

We subtracted the distance along the actin contour from the pointed end (defined as the abscissa in Cytosim) at the end of each binding event from the distance from the pointed end at the beginning to analyze the displacement per binding event in Figure S2 B. We calculated the mean force on the myosin-actin bond among all time points in the lifetime of a binding event for the mean force measurement in Figure S2 C.

For individual binding event analyses in Figure S4 C-E, we calculated the difference in the given actin measurement (barbed end localization or angle) from the simulation time point immediately before each binding event to the first time point of the binding event. We performed the same calculation immediately before and after the end of the bond lifetime for unbinding events as well.

#### 0.0.7 Actin density and localization analysis

Density of assembled actin subunits at an endocytic site was calculated as the total count of actin model points within a given volume. The density for Figure S5 A was calculated for the volume of the whole simulation system. The density calculated for Figure S4 A-B was filtered just for actin barbed ends, within the volume of each radial distance and z-distance bin. The count of daughter filaments for Figure S5 B is analogous to the density of bound Arp2/3 complexes in the branched actin network. Filament length for Figure S5 C was calculated as the contour length for actin filaments in the simulation. Elongation attenuation for Figure S5 D was defined as the difference between measured per-filament elongation rate and the theoretical maximum elongation rate. All measurements for Figure S5 were filtered for filaments either directly bound to Hip1R linkers or indirectly bound through mother or daughter filaments.

#### 0.0.8 Bending energy analysis

Per-filament actin bending energy for Figure S6 C was calculated from the difference in tangential direction between the barbed end and the closest Hip1R attachment by contour length. Only filaments that were directly or indirectly bound to Hip1R were included in this analysis.

## Supplementary Information

**Table S1:**
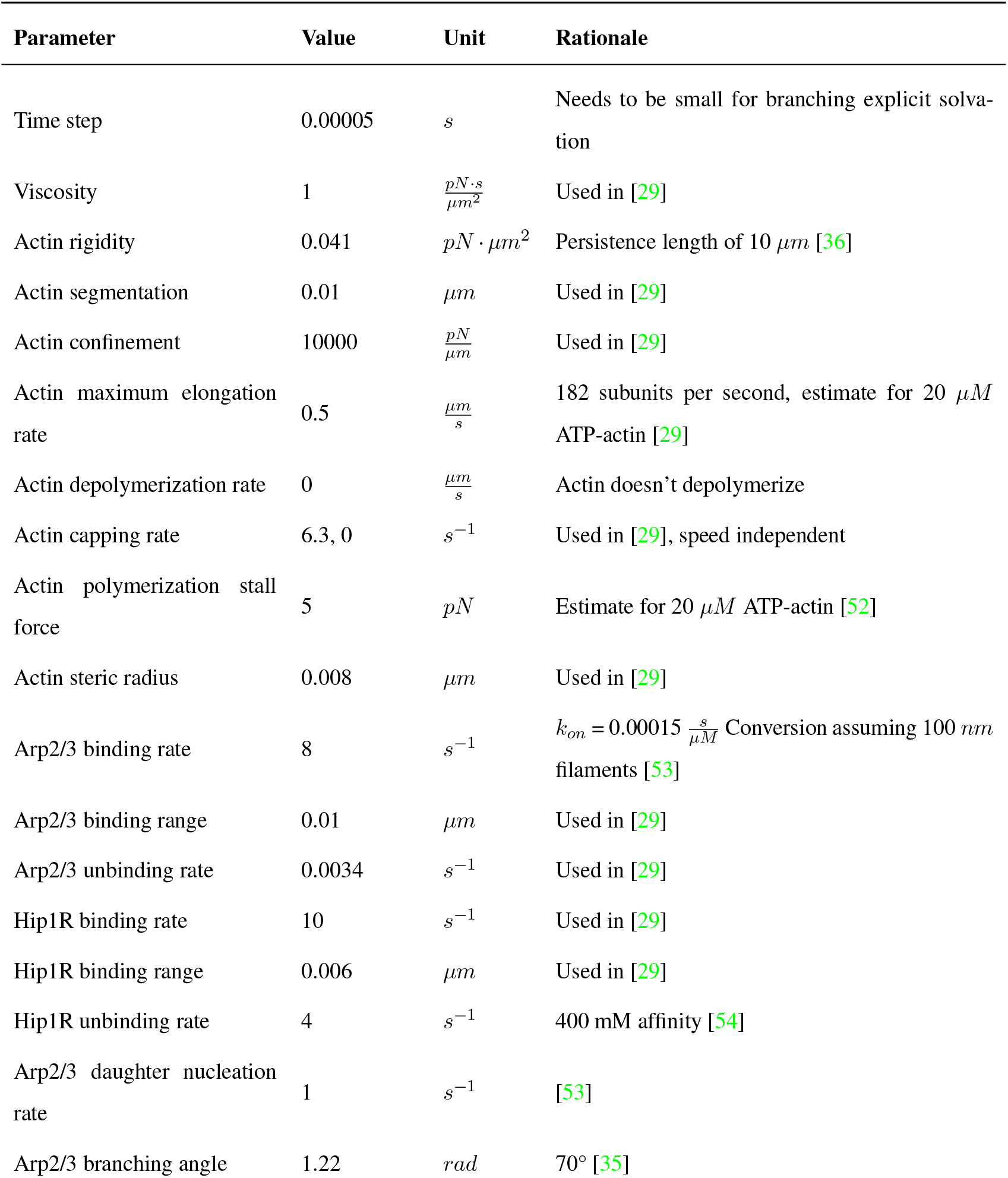
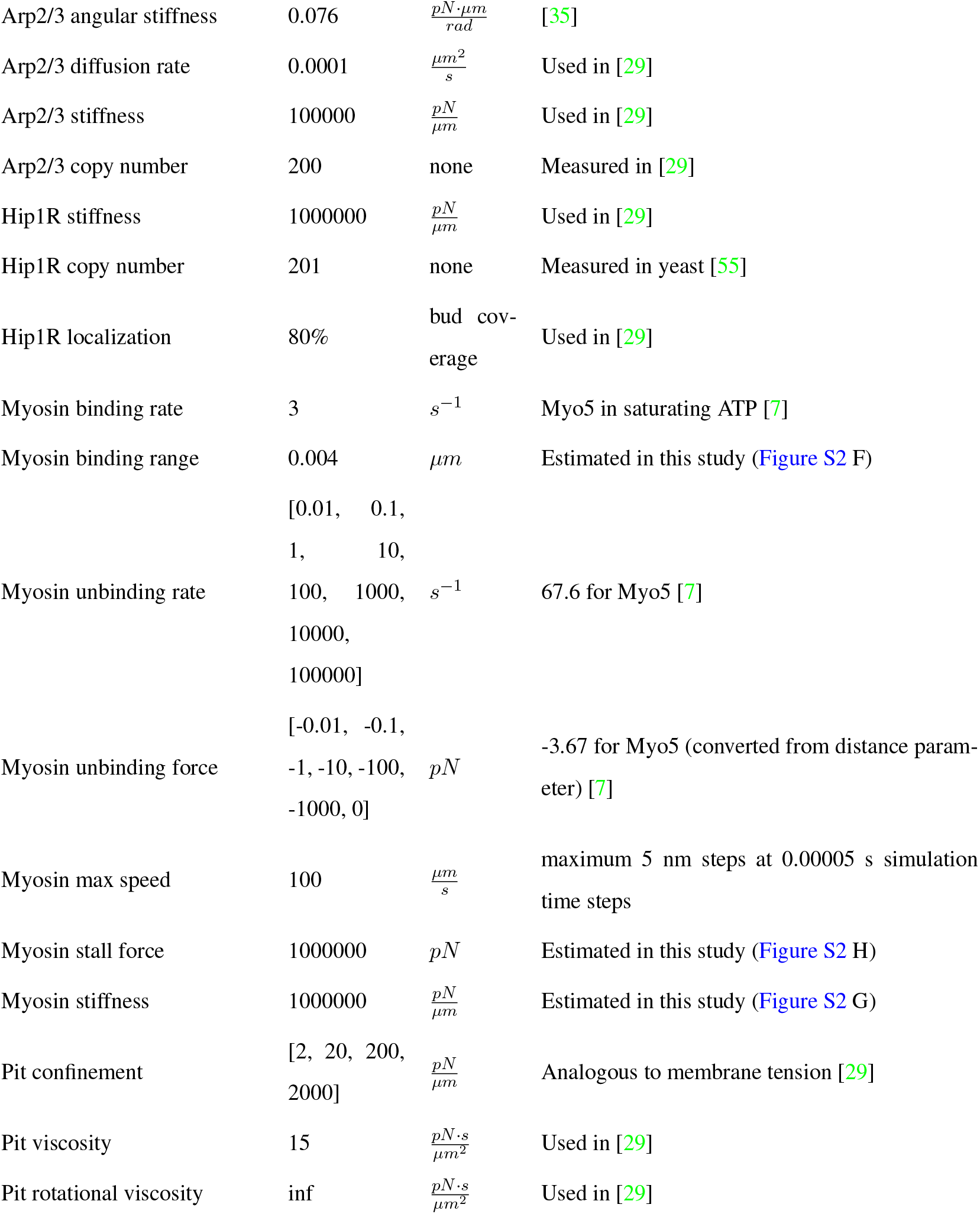
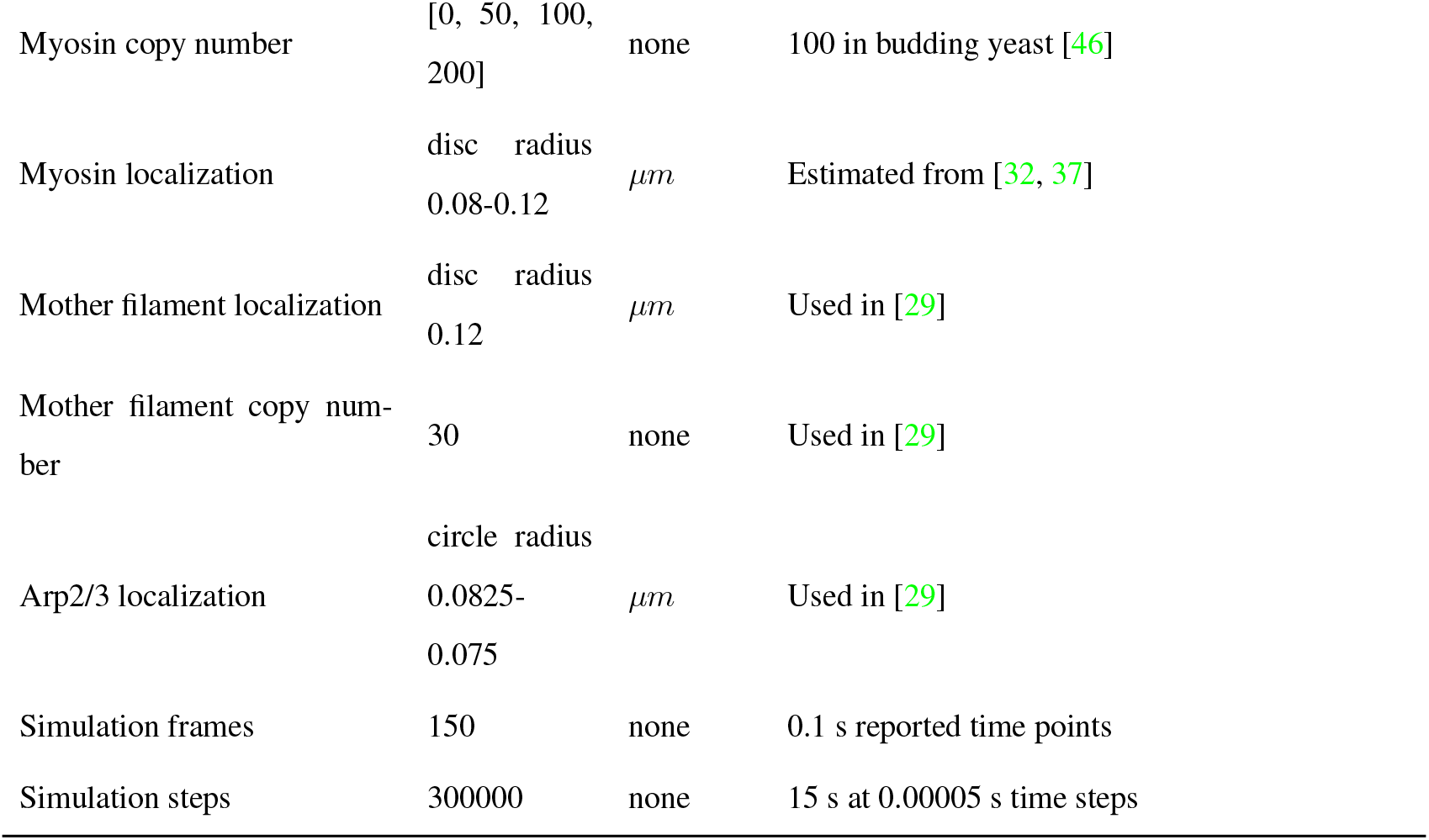
Model Parameters.

| Parameter | Value | Unit | Rationale |
| --- | --- | --- | --- |
| Time step | 0.00005 | $s$ | Needs to be small for branching explicit solvation |
| Viscosity | 1 | $\frac{pN \cdot s}{\mu m^2}$ | Used in [29] |
| Actin rigidity | 0.041 | $pN \cdot \mu m^2$ | Persistence length of 10 $\mu m$ [36] |
| Actin segmentation | 0.01 | $\mu m$ | Used in [29] |
| Actin confinement | 10000 | $\frac{pN}{\mu m}$ | Used in [29] |
| Actin maximum elongation rate | 0.5 | $\frac{\mu m}{s}$ | 182 subunits per second, estimate for 20 $\mu M$ ATP-actin [29] |
| Actin depolymerization rate | 0 | $\frac{\mu m}{s}$ | Actin doesn't depolymerize |
| Actin capping rate | 6.3, 0 | $s^{-1}$ | Used in [29], speed independent |
| Actin polymerization stall force | 5 | $pN$ | Estimate for 20 $\mu M$ ATP-actin [52] |
| Actin steric radius | 0.008 | $\mu m$ | Used in [29] |
| Arp2/3 binding rate | 8 | $s^{-1}$ | $k_{on} = 0.00015 \frac{s}{\mu M}$ Conversion assuming 100 $nm$ filaments [53] |
| Arp2/3 binding range | 0.01 | $\mu m$ | Used in [29] |
| Arp2/3 unbinding rate | 0.0034 | $s^{-1}$ | Used in [29] |
| Hip1R binding rate | 10 | $s^{-1}$ | Used in [29] |
| Hip1R binding range | 0.006 | $\mu m$ | Used in [29] |
| Hip1R unbinding rate | 4 | $s^{-1}$ | 400 mM affinity [54] |
| Arp2/3 daughter nucleation rate | 1 | $s^{-1}$ | [53] |
| Arp2/3 branching angle | 1.22 | $rad$ | 70° [35] |
| Arp2/3 angular stiffness | 0.076 | $\frac{pN \cdot \mu m}{rad}$ | [35] |
| Arp2/3 diffusion rate | 0.0001 | $\frac{\mu m^2}{s}$ | Used in [29] |
| Arp2/3 stiffness | 100000 | $\frac{pN}{\mu m}$ | Used in [29] |
| Arp2/3 copy number | 200 | none | Measured in [29] |
| Hip1R stiffness | 1000000 | $\frac{pN}{\mu m}$ | Used in [29] |
| Hip1R copy number | 201 | none | Measured in yeast [55] |
| Hip1R localization | 80% | bud coverage | Used in [29] |
| Myosin binding rate | 3 | $s^{-1}$ | Myo5 in saturating ATP [7] |
| Myosin binding range | 0.004 | $\mu m$ | Estimated in this study (Figure S2 F) |
| Myosin unbinding rate | [0.01, 0.1, 1, 10, 100, 1000, 10000, 100000] | $s^{-1}$ | 67.6 for Myo5 [7] |
| Myosin unbinding force | [-0.01, -0.1, -1, -10, -100, -1000, 0] | $pN$ | -3.67 for Myo5 (converted from distance parameter) [7] |
| Myosin max speed | 100 | $\frac{\mu m}{s}$ | maximum 5 nm steps at 0.00005 s simulation time steps |
| Myosin stall force | 1000000 | $pN$ | Estimated in this study (Figure S2 H) |
| Myosin stiffness | 1000000 | $\frac{pN}{\mu m}$ | Estimated in this study (Figure S2 G) |
| Pit confinement | [2, 20, 200, 2000] | $\frac{pN}{\mu m}$ | Analogous to membrane tension [29] |
| Pit viscosity | 15 | $\frac{pN \cdot s}{\mu m^2}$ | Used in [29] |
| Pit rotational viscosity | inf | $\frac{pN \cdot s}{\mu m^2}$ | Used in [29] |
| Myosin copy number | [0, 50, 100, 200] | none | 100 in budding yeast [46] |
| Myosin localization | disc radius<br>0.08-0.12 | $\mu m$ | Estimated from [32, 37] |
| Mother filament localization | disc radius<br>0.12 | $\mu m$ | Used in [29] |
| Mother filament copy number | 30 | none | Used in [29] |
| Arp2/3 localization | circle radius<br>0.0825-0.075 | $\mu m$ | Used in [29] |
| Simulation frames | 150 | none | 0.1 s reported time points |
| Simulation steps | 300000 | none | 15 s at 0.00005 s time steps |

**Figure S1:**
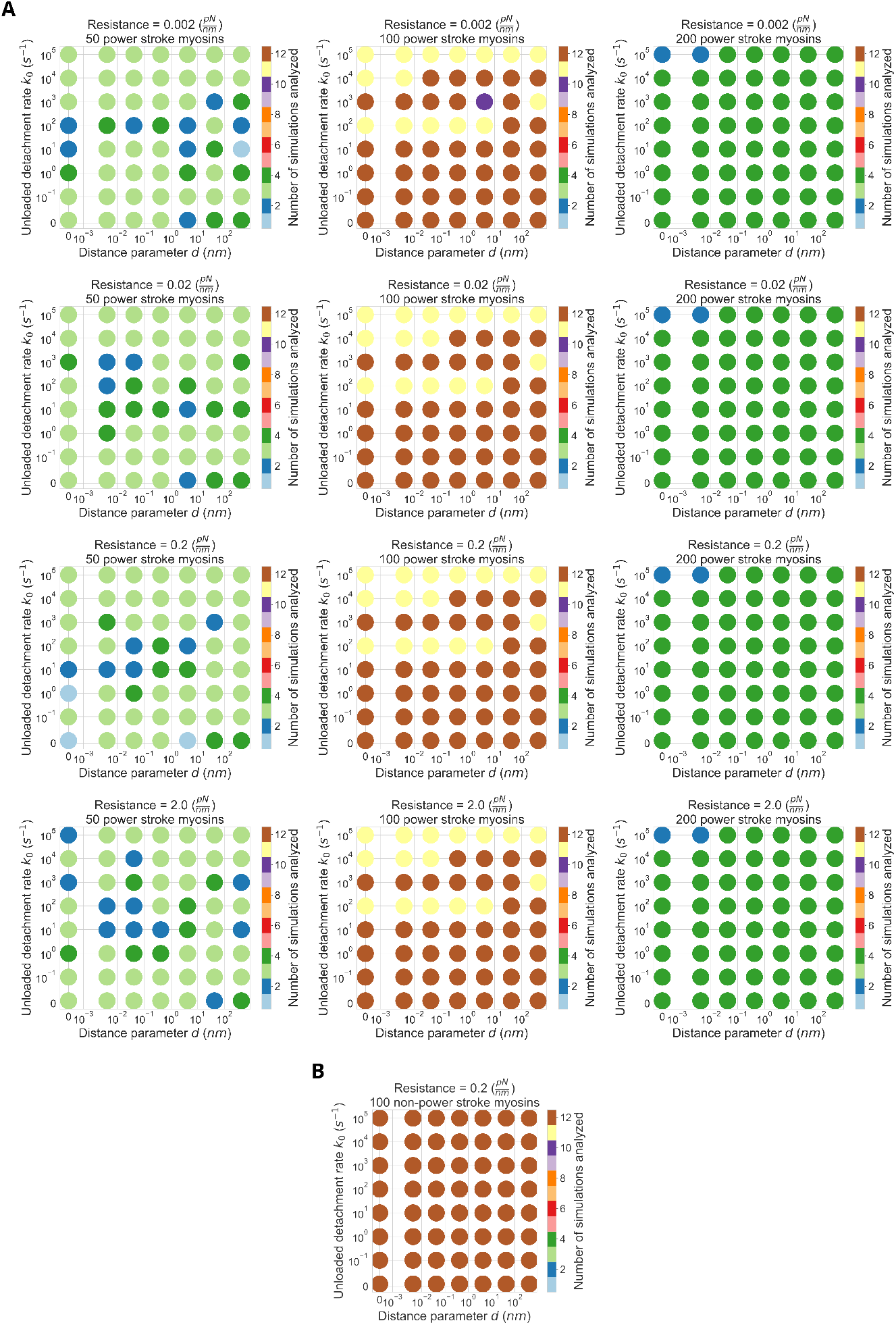
Number of replicate simulations ran to completion for analysis in this study with (A) and without (B) motor activity.

**Figure S2:**
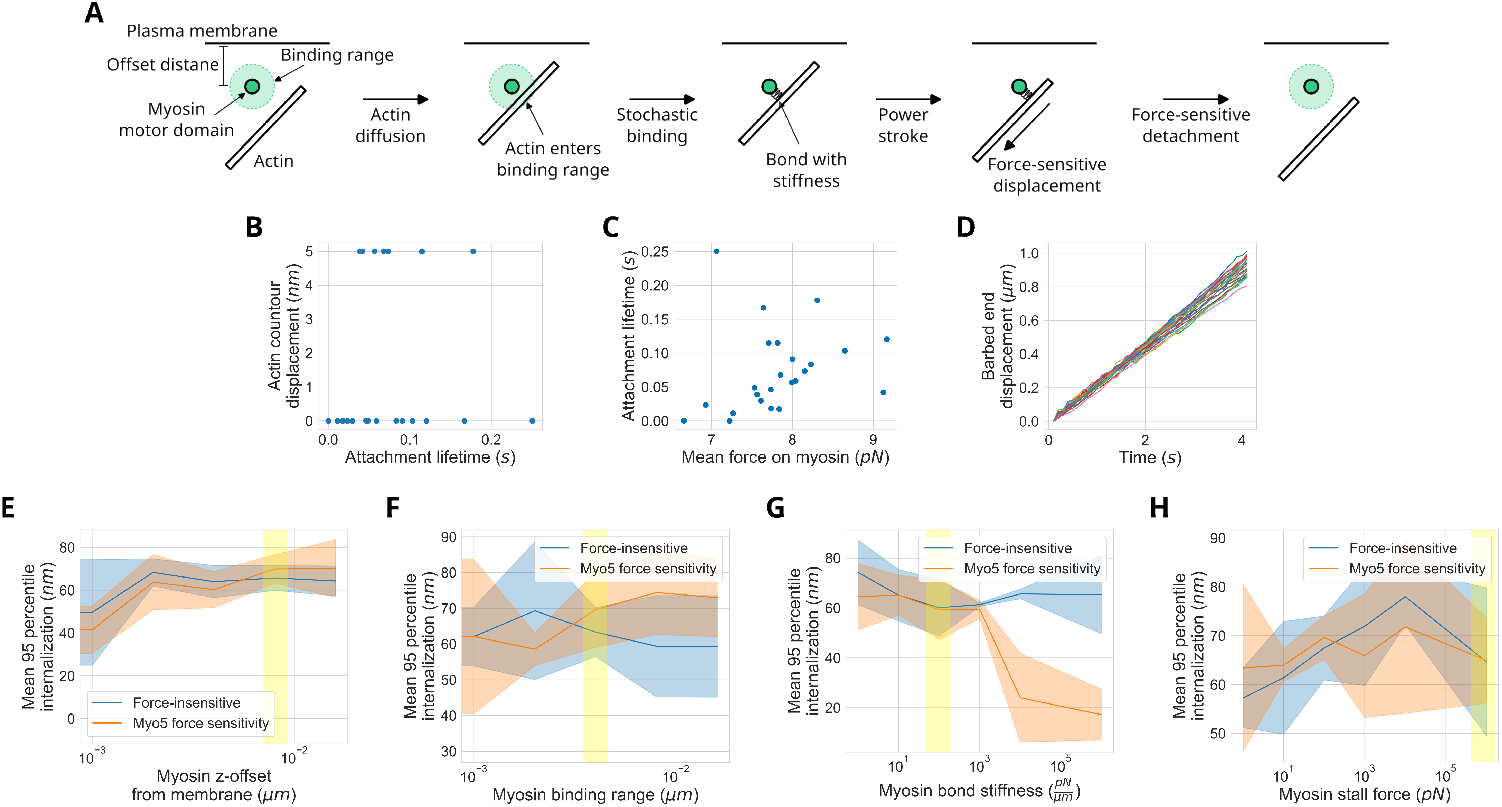
Design of a model Myosin-I in Cytosim. (A) Cartoon depiction of Myosin-I activity in simulations. A myosin is modeled as a single motor domain at a fixed offset distance from the plasma membrane base. When an actin filament diffuses into the binding range of the myosin, it can form a bond at a probability set by the binding rate parameter. Like all bonds in Cytosim, the actin-myosin bond behaves like a spring with a set stiffness. Immediately after forming the bond, the myosin exerts a force on the actin to displace it up to 5 nm, depending on the applied force and myosin’s stall force. Following this power stroke, the bond remains until it unbinds at a probability set by the unloaded detachment rate and distance parmameter (Equation 1). (B) Validation of non-processive motor activity. In contrast to the default motor activity in Cytosim in which displacement increases linearly with time, our model myosin-I only displaces up to 5 nm in a single step per attachment event. (C) Validation of force-sensitive detachment. Similarly to single-molecule measurements of Myo5 [50], the lifetime of simulated myosin-actin attachment events generally increases as the mean applied force on the bond increases. (D) Functional validation of model type I myosins in a simulated 2D gliding assay. The displacement of individual actin filaments gliding over a substrate coated in myosins are plotted over time. The traces show actin velocities of 0.2-0.25 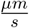, compared to 0.8 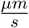 measured in the analogous experiment for Myo5 [50]. (E-H) Parameter sensitivity analysis for experimentally unconstrained parameters. The mean 95th percentile of internalization with error bars for standard deviation (n=4) were plotted for CME simulations with either myosins with distance parameter of 0 (blue) or with Myo5-constrained distance parameter (orange) [50]. The values used for all other simulations in this study are highlighted in yellow. All simulations were run with a resistance to pit internalization of 0.2 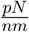.

**Figure S3:**
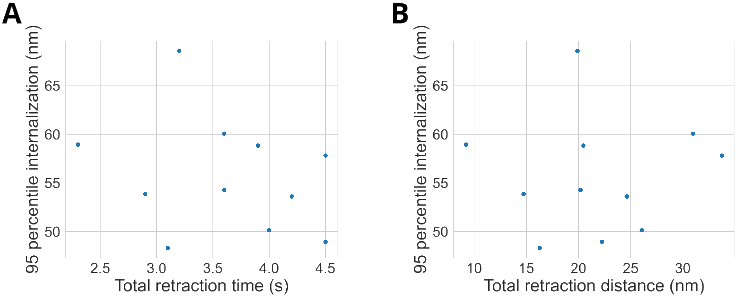
Pit retraction measurements (A) total retraction time and (B) total retraction distance do not correlate with overall internalization in the absence of myosin. Points represent individual replicate simulations where no myosins were included. All simulations were run with a resistance to pit internalization of 0.2 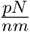.

**Figure S4:**
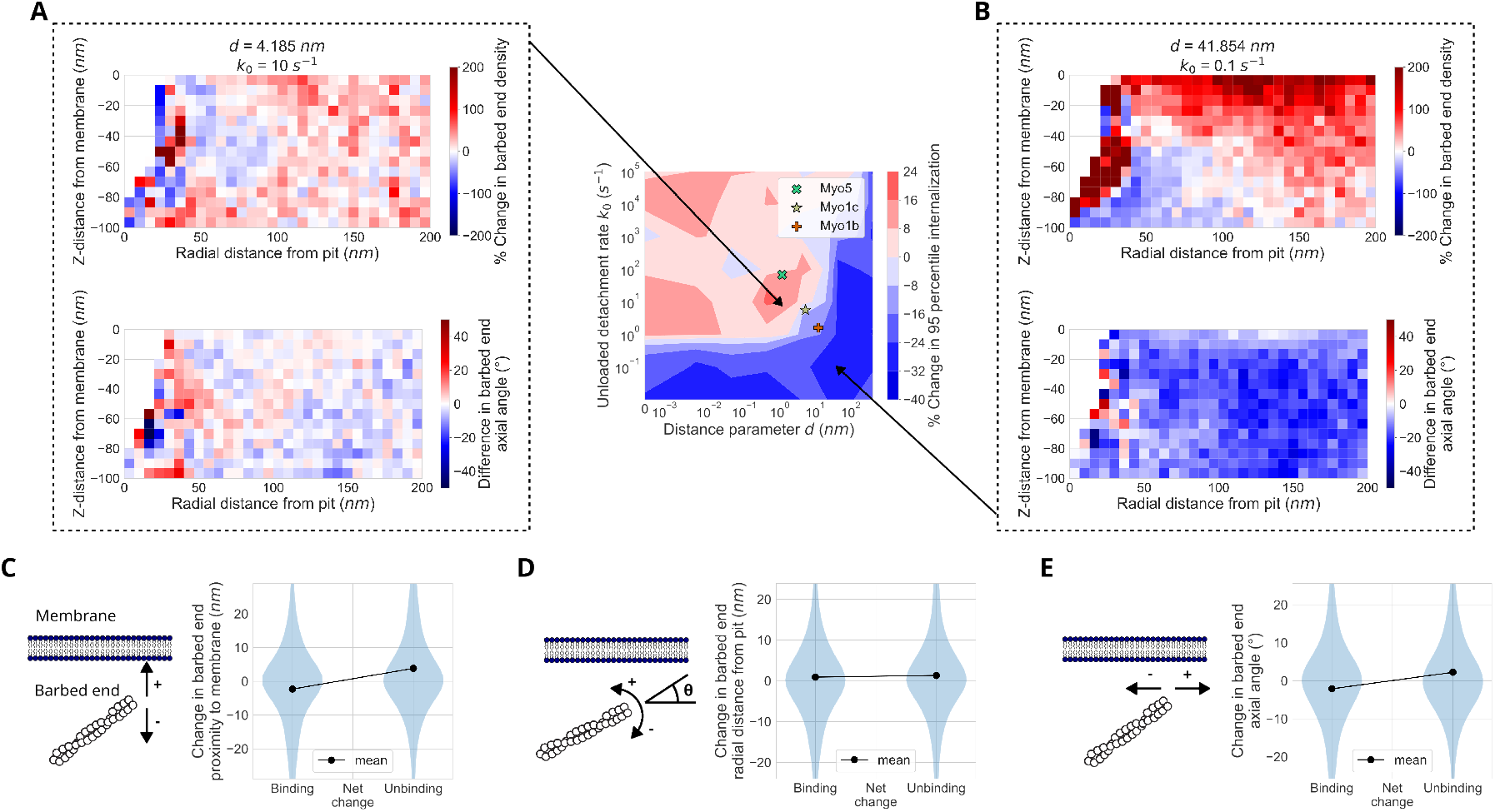
Myosin-I subtly reorganizes the actin network to assist internalization. (A) % change in mean barbed end localization (top) and angle relative to the plasma membrane (bottom, diagrammed in D) relative to simulations with 0 myosins, over radial distance from the endocytic pit and z-distance from the membrane, for myosins with assistive kinetic parmaters indicated by the arrow to the middle plot. Note the relative enrichment of barbed ends near the membrane at 75-200 nm radial distance from the pit. (B) Changes in barbed end localization and angle with the plasma membrane for myosins with internalization-inhibiting kinetic parameters indicated by the arrow to the middle plot. (C-E) Changes to actin barbed end z-position relative to the membrane (C), angle relative to the membrane (D), and radial distance from the pit (E) immediately after binding and unbinding myosin. Each plot represents the distribution of all binding/unbinding events in the last 5 seconds of simulations with myosins with assistive kinetic parameters (the same simulations analyzed in A). All simulations were run with a resistance to pit internalization of 0.2 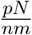.

**Figure S5:**
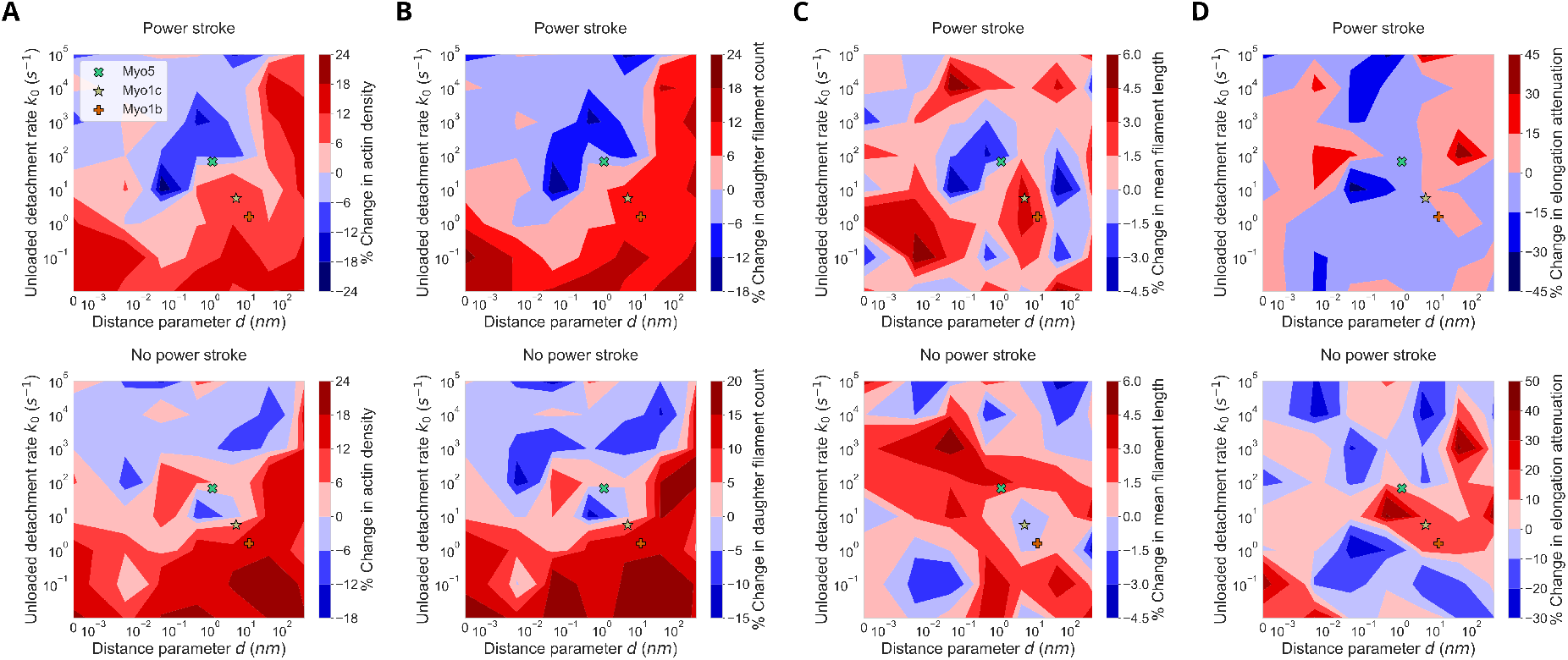
Myosin-I does not assist internalization through increasing actin assembly. (A) Contour plots of the % change in mean total hip1r-associated actin monomers in the last 5 seconds of simulations relative to simulations with 0 myosins, over a range of parameter values for *d* and *k*_0_. At low *d* and high *k*_0_, myosins cause a slight decrease in actin accumulation, while they cause an increase in actin accumulation at high *d* and low *k*_0_. There is a subtle shift in the boundary between increasing and decreasing actin accumulation in simulations with myosins that have motor stepping enabled (top) compared to disabled (bottom). (B) Contour plot of the % change in mean total hip1r-associated actin filaments in the last 5 seconds of simulations relative to simulations with 0 myosins. The nearly identical pattern to (A) indicates that the changes in total mass of actin are explained by changes in daughter filament nucleation. (C) Contour plot of the % change in mean length of hip1r-associated actin filaments in the last 5 seconds of simulations relative to simulations with 0 myosins. There is no discernable pattern in the myosin parameters, regardless of motor activity. (D) Contour plot of the % change in elongation attenuation of hip1r-associated actin filaments in the last 5 seconds of simulations relative to simulations with 0 myosins. There is no discernable pattern in the myosin parameters, regardless of motor activity. All simulations were run with a resistance to pit internalization of 0.2 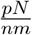.

**Figure S6:**
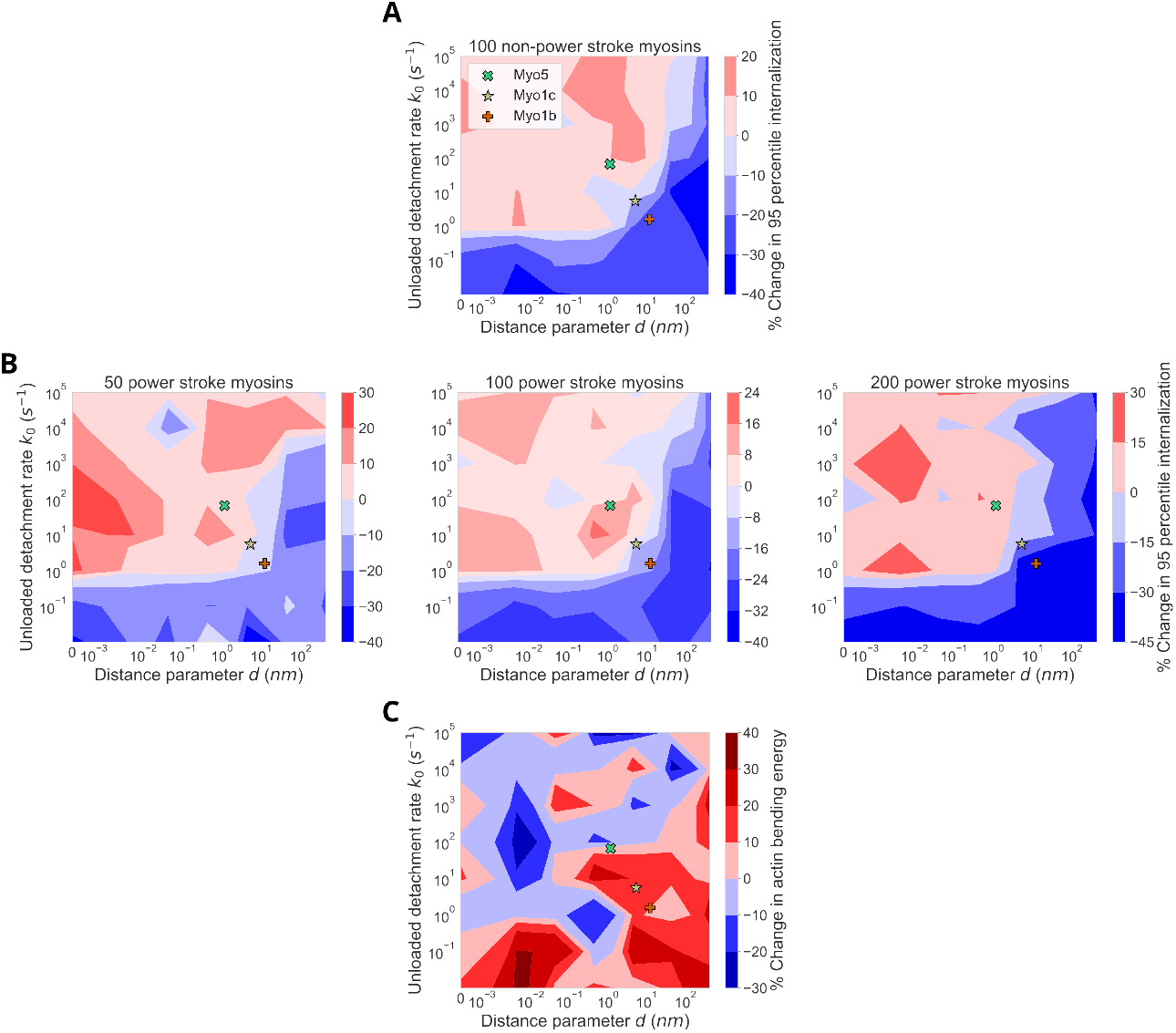
Myosin-I does not assist internalization through motor force transduction. (A) Contour plot of the % change in mean 95th percentile of internalization for simulations with myosin motor activity disabled relative to simulations with 0 myosins, over a range of parameter values for *d* and *k*_0_. The boundary lies at a very similar location to that of simulations with motor activity (B, middle). (B) No clear dose-dependence of myosins on internalization. Countour maps of 95th percentile internalization for simulations with 50 (left), 100 (middle), and 200 (right) myosins, relative to 0 myosins. (C) Contour plot of the % change in mean bending energy of hip1r-bound actin filaments in the last 5 seconds of simulations relative to simulations with 0 myosins. An increase in bending energy is not associated with increased internalization. All simulations were run with a resistance to pit internalization of 0.2 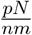.

